# Pharmaceutical pollution alters the cost of bacterial infection and its relationship to pathogen load

**DOI:** 10.1101/2023.08.22.554372

**Authors:** Lucinda C. Aulsebrook, Bob B.M. Wong, Matthew D. Hall

## Abstract

The relationship between pathogen proliferation and the cost of infection experienced by a host drives the ecology and evolution of host-pathogen dynamics. While environmental factors can shape this relationship, there is currently limited knowledge on the consequences of emerging contaminants, such as pharmaceutical pollutants, for the commonly observed trade-off between a pathogen’s growth within the host and the damage it causes, termed its virulence. Here, we investigated how exposure to fluoxetine (Prozac), a commonly detected psychoactive pollutant, could alter this key relationship using the water flea *Daphnia magna* and its bacterial pathogen *Pasteuria ramosa* as a model system. Across a variety of fluoxetine concentrations, we found that fluoxetine shaped the damage a pathogen caused, such as the reduction in fecundity or intrinsic growth experienced by infected individuals, but with minimal change in average pathogen spore loads. Instead, fluoxetine modified the relationship between the degree of pathogen proliferation and its virulence, with both the strength of this trade-off and the component of host fitness most affected varying by fluoxetine concentration and host genotype. Our study underscores the potential for pharmaceutical pollution to modify the virulence of an invading pathogen, as well as the fundamental trade-off between host and pathogen fitness, even at the trace amounts increasingly found in natural waterways.

## Introduction

Pathogens are ubiquitous across ecosystems, and the harm they inflict on their hosts has important ramifications for both host and pathogen ecology and evolution [1, 2]. The fitness costs that a pathogen inflicts on its host upon infection, such as increased mortality or reduced fecundity, is referred to as the pathogen’s virulence [1, 3], and is seen as an unavoidable consequence of pathogens exploiting host resources to increase their own transmission success [3, 4]. Both host and pathogen evolution, however, can shape this relationship [1, 3]. For a pathogen, increased virulence must be balanced against the potential transmission costs of a shorter infectious period or reduced overall proliferation that arises if a host dies too early [3-5]. In turn, hosts can evolve to fight infection by either reducing pathogen burden via limiting pathogen growth (i.e., resistance), or by minimising the damage a pathogen causes (i.e., tolerance) [6-8].

This relationship between virulence and the proliferation of a pathogen within a host is integral to theory surrounding disease dynamics [2, 9], and has been shown to depend on a variety of biotic and abiotic factors, including host and pathogen genotype [10], the sex and age of the host [11], the transmission mode of the pathogen [12], as well as the environment [13, 14]. In particular, the environmental stressors that host organisms confront often reduce their condition, which can result in the expression of higher virulence upon infection [15]. Furthermore, environmental stressors may increase the virulence of a pathogen via reducing the resistance or tolerance of a host, due to the energetic cost involved in coping with these stressors [16, 17]. For example, pollutants such as pesticides and heavy metals have been shown to result in increased virulence of various parasites in a range of hosts [13, 18-21]. Such increases in virulence are often predicted to result in a reduction in transmission due to earlier host mortality [18], although corresponding changes in pathogen loads are not always found (e.g. [21]). Whether pollutants induce changes in the transmission-virulence relationship via changes in virulence, pathogen replication, or both, is central to understanding the role pollution plays in shaping host-pathogen interactions.

An environmental stressor of increasing concern is pharmaceutical pollution, with hundreds of products being detected globally in rivers, lakes and waterways [22, 23]. Following usage, pharmaceuticals are often excreted still in a bioactive form, which leads to the release of these compounds into the environment via wastewater outlets [24]. These contaminants are then slow to degrade in the environment, and can bioaccumulate [25], with concerning implications for exposed wildlife [26], such as feminisation of male fish [27-29] and renal failure in vultures [30]. However, little is known about how these pharmaceutical pollutants influence infectious disease traits (see [31, 32]), such as the cost of infection or transmission potential, and thus how pharmaceuticals may affect this trade-off, which is fundamental to disease ecology and evolution.

While pollutants such as heavy metals and pesticides frequently decrease host condition or reproductive output [33-35], exposure to pharmaceutical pollutants have sometimes been reported to induce positive effects on growth and reproduction [36, 37]. Such effects could potentially result in decreasing the fitness costs inflicted by a pathogen, which could, in turn, have ramifications for pathogen replication. Unlike legacy pollutants (e.g. pesticides, heavy metals), the effects of pharmaceuticals are also frequently non-monotonic, where lower concentrations can result in more severe responses than higher concentrations (e.g. [36, 38, 39]), perhaps due to these drugs being designed to act in a selective dose-dependent manner [40]. As a result, even trace amounts of pharmaceutical pollutants in the environment have the potential to impact ecosystem dynamics, possibly to a greater degree than areas subject to stronger pollution. These characteristics may result in pharmaceutical pollutants affecting pathogen virulence and transmission in a manner that is quite distinct from more traditional pollutants.

One concerning pharmaceutical contaminant is fluoxetine, marketed as Prozac. Fluoxetine is one of the most commonly prescribed antidepressants globally [41, 42], as well as one of the most frequently detected psychoactive pollutants in the environment [25, 43]. As a selective serotonin reuptake inhibiter, fluoxetine acts to prevent the reuptake of the neurotransmitter serotonin by interacting with the serotonin transporter (5-HTT or SERT), causing the effect of serotonin to be prolonged [44]. Since serotonin and its transporters are evolutionary conserved in a wide range of taxa, it is unsurprising that exposure to fluoxetine has been found to have effects in a variety of wildlife, including non-monotonic effects on reproduction in invertebrates (e.g. [36, 45]) and behaviour in numerous fish species (e.g. [39, 46, 47]). However, almost nothing is known about how environmental levels of psychoactive pollutants such as fluoxetine could influence the cost of infection to hosts or transmission potential of pathogens (but see [31]), and thus how these contaminants may shape the ecology and evolution of host-parasite dynamics.

In our study, we investigated the consequences of fluoxetine exposure on the cost of infection and pathogen replication, using *Daphnia magna* and its bacterial pathogen *Pasteuria ramosa*. We chronically exposed *D. magna* to two environmentally realistic concentrations of fluoxetine (i.e., nominal concentrations of 30ng/L and 300ng/L), as well as a concentration used for acute toxicology tests (3000ng/L), and a freshwater control. We then measured the reduction in host fitness factors, such as fecundity and intrinsic growth, to obtain an indication of virulence. Additionally, we recorded the spore loads of infected individuals as a measure of pathogen replication, which is commonly used as a proxy for transmission in many study systems [9]. By examining how fluoxetine exposure shifts the relationship between the cost of infection and pathogen replication, we explored whether this fundamental trade-off is sensitive to pharmaceutical pollution. Such information will assist predictions of how pharmaceutical pollution may uniquely shape the evolution and ecology of host-pathogen interactions.

## Methods

### Study System

*Daphnia magna* is a freshwater filter-feeding crustacean that is frequently used as a model in aquatic toxicology (e.g. [48]), as they are sensitive to their environment and provide an essential role in aquatic ecosystems as primary consumers [49]. *Daphnia magna* and the bacterial pathogen *P. ramosa* are also commonly used in disease ecology studies, due to the influence of environmental factors on both host and pathogen fitness (reviewed in [50]). *Daphnia magna* become infected with *P. ramosa* via ingesting spores present in water or sediment [50]. If not cleared in the first few days, the infection is chronic and results in severe reduction in host fecundity, and an increase in body size [50-53]. Upon the death of the *D. magna, P. ramosa* spores are released from the cadaver, whereupon they may be ingested by other *D. magna*.

For this study, we used two *D. magna* genotypes derived from single clones: HU-HO-2 (herein HO2) from Hungary and BE-OHZ-M10 (herein M10) from Belgium. These genotypes were chosen as they are known to differ in several life-history traits as well as infection outcomes (e.g. [51, 54]), allowing insights into whether fluoxetine effects are likely to be genotype specific. For three generations prior to the experiment, animals from each clone were cultured individually in 70mL jars, filled with 45mL of artificial *Daphnia* media [55, 56], which was replaced twice a week. Algae (*Scenedesmus spp*.) was dispensed into each jar daily according to the growing needs of the animals, from 0.5 million cells per animal on day 1, to 5 million cells per animal from day 8 onwards. All animals were kept in a controlled-temperature room with a constant temperature of 20°C and an 18:6h light dark cycle. Experimental animals were taken from the third or fourth clutch of parental *D. magna* and maintained under the same standard conditions as parental generations.

### Fluoxetine and pathogen exposure

From day 1, *D. magna* were exposed chronically to one of the four nominal fluoxetine concentrations: low (30ng/L), medium (300ng/L), high (3000ng/L), and a freshwater control (0ng/L). The low and medium concentrations are within the range that has previously been detected in the environment, where 30ng/L represents levels detected in surface water and 300ng/L represents levels found in direct effluent flows from wastewater treatment plants [57]. The high treatment represents a nominal concentration that is less environmentally realistic but represents the magnitude used in acute toxicology studies [58]. These fluoxetine treatments were produced following established protocols [36, 39, 59, 60], by dissolving fluoxetine hydrochloride in a small volume of methanol, and then distributing this methanol into the *Daphnia* media. The control treatment was dosed with the same volume of methanol, but contained no fluoxetine. Fresh dosing of fluoxetine occurred twice weekly at each water change.

Weekly water samples from each fluoxetine treatment were collected after dosing, and sent to Envirolab Services (MPL Laboratories; NATA accreditation: 2901; accredited for compliance with ISO/IEC: 17025) for analysis. Fluoxetine concentrations were derived using gas chromatography-tandem mass spectrometry (7000C Triple Quadrupole GC-MS/MS, Agilent Technologies, Delaware, USA) following methods described in [39]. Measured concentrations were in line with nominal fluoxetine doses (low: 25.87 ± 2.47ng/L, medium: 197.5 ± 10.3ng/L and high: 1900 ± 184ng/L, see supplementary material), capturing a tenfold difference in concentration between each treatment.

At day 4 and 5, *D. magna* in the disease exposure treatment received 20,000 *P. ramosa* spores (C1 genotype). Our study utilised a full factorial design with 20-36 individuals per treatment (2 *D. magna* genotypes × 2 disease treatments (exposed or unexposed) × 4 fluoxetine treatments). Variation in the number of individuals per treatment was due to differences in infection rate, as well as handling errors. Throughout the experiment, *D. magna* were monitored daily for survival, and dead individuals were frozen in 0.5mL RO water at -20°C. Offspring were counted at each water change. At 30 days, the body size of all remaining *D. magna* was measured using a scaled binocular microscope and all *D. magna* were frozen individually in 0.5mL RO water for later inspection of spore production. Intrinsic growth rates (r) were calculated using the timing and number of offspring and then solving the Euler-Lotka equation (following [36, 61]).

### Spore analysis

Spore counts per animal were measured using an Accuri C6 flow cytometer (BD Bioscience, San Jose, CA) as per [54]. Infected individuals previously frozen in 0.5mL RO water were thawed and crushed, and 10uL of each sample was transferred into 190μL 5mM EDTA in a round-bottomed PPE 96-well plate. Each run involved counting 32 wells, which included a total of 12 *D. magna* individuals, each counted twice, as well as 8 wells containing EDTA only as a wash step. Mature spores were identified and quantified based on their distinct size, morphology and fluorescence (in contrast to immature spores, algae or animal debris) using gates base on fluorescence (via the 670 LP filter) and side scatter (cell granularity).

### Statistical analysis

Statistical tests were performed using R software version 4.0.3 software (R Development Core Team, 2020). First, we estimated the costs of infection, in regards to the relative changes in fecundity, body size and intrinsic growth, by calculating the difference between each individual’s trait value and the mean of the corresponding uninfected animals of the same genotype and fluoxetine treatment. We then analysed changes in these cost of infection traits (reduction in fecundity, reduction in intrinsic growth, increase in body size) as well as spore loads, via a linear mixed effects model (via lme4 [62]) with fluoxetine treatment, disease exposure, genotype and interactive terms as fixed effect factors, and block as a random effect. Residuals from all fitted models were approximately normally distributed. Post-hoc tests were generated using the ‘emmeans’ package [63] to produce pair-wise comparisons.

We then explored how fluoxetine may also change the relationship between a pathogen’s spore load and the fitness costs it imposes on a host, by fitting a series of multiple regression models to the data, akin to an extension of a two-factor analysis of covariance. Each model considered the linear relationships between spore load (the response variable) and the three costs of infection traits (the regression coefficients), with the host and fluoxetine treatments as additional fixed effects and blocks as a random effect. The series of models tested specific hypotheses regarding how each treatment potentially influenced the partial effects of fluoxetine on spore loads, ranging from a single pattern for all treatments (no interaction terms included, regression coefficients the same for all treatments), to separate patterns for every combination of host genotype and fluoxetine level (including three-way interaction terms, regression coefficients varying in all treatments). Each model containing the terms of interest was compared to a reduced model and the improvement in fit evaluated using a log-likelihood ratio test. Before analysis, in each treatment we rescaled pathogen spore loads to a mean of one (i.e., relative spore loads) and standardised the fitness cost measures (mean = 1, standard deviation = 0), following standard selection analysis approaches [64].

## Results

### Fluoxetine exposure did not alter pathogen replication, but shifted the cost of infection

We found that the number of spores produced per individual did not appear to be significantly influenced by fluoxetine exposure for either host genotype, indicating no change in pathogen replication across a 100-fold change in fluoxetine concentration (Table 1, Figure 1A). There were also no significant differences in survival between fluoxetine treatment groups within the 30-day period. Conversely, the cost of infection, in terms of changes in fecundity, intrinsic growth and body size, appeared to differ between fluoxetine treatment groups. The relative decrease in fecundity experienced by an infected individual was dependent on an interaction between fluoxetine treatment and genotype (Table 1). For both host genotypes, infected *D. magna* exposed to the medium and high fluoxetine concentrations experienced a greater reduction in the number of offspring produced compared to the control treatment. However, for M10 only, infected *D. magna* exposed to the low fluoxetine concentration had a smaller decrease in offspring than the control (i.e., freshwater) group (Figure 1B).

**Table 1.**
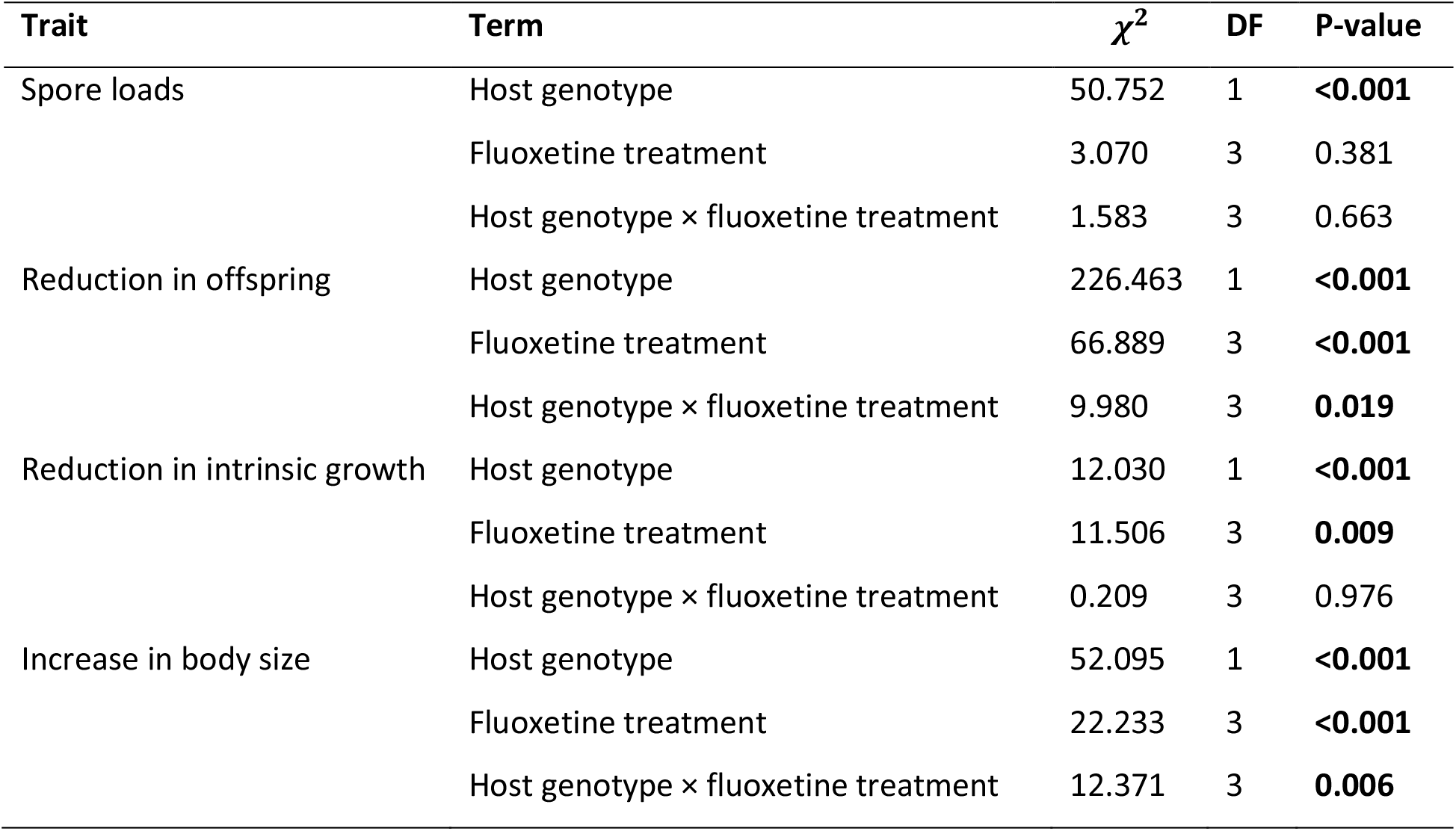
Effects of host genotype, fluoxetine, and their interaction on spore loads, reduction in offspring, reduction in intrinsic growth and increase in body size of infected *Daphnia magna*. Analysis was performed using liner mixed effect models.

| Trait | Term | $\chi^2$ | DF | P-value |
| --- | --- | --- | --- | --- |
| Spore loads | Host genotype | 50.752 | 1 | **<0.001** |
|  | Fluoxetine treatment | 3.070 | 3 | 0.381 |
|  | Host genotype × fluoxetine treatment | 1.583 | 3 | 0.663 |
| Reduction in offspring | Host genotype | 226.463 | 1 | **<0.001** |
|  | Fluoxetine treatment | 66.889 | 3 | **<0.001** |
|  | Host genotype × fluoxetine treatment | 9.980 | 3 | **0.019** |
| Reduction in intrinsic growth | Host genotype | 12.030 | 1 | **<0.001** |
|  | Fluoxetine treatment | 11.506 | 3 | **0.009** |
|  | Host genotype × fluoxetine treatment | 0.209 | 3 | 0.976 |
| Increase in body size | Host genotype | 52.095 | 1 | **<0.001** |
|  | Fluoxetine treatment | 22.233 | 3 | **<0.001** |
|  | Host genotype × fluoxetine treatment | 12.371 | 3 | **0.006** |

**Figure 1.**
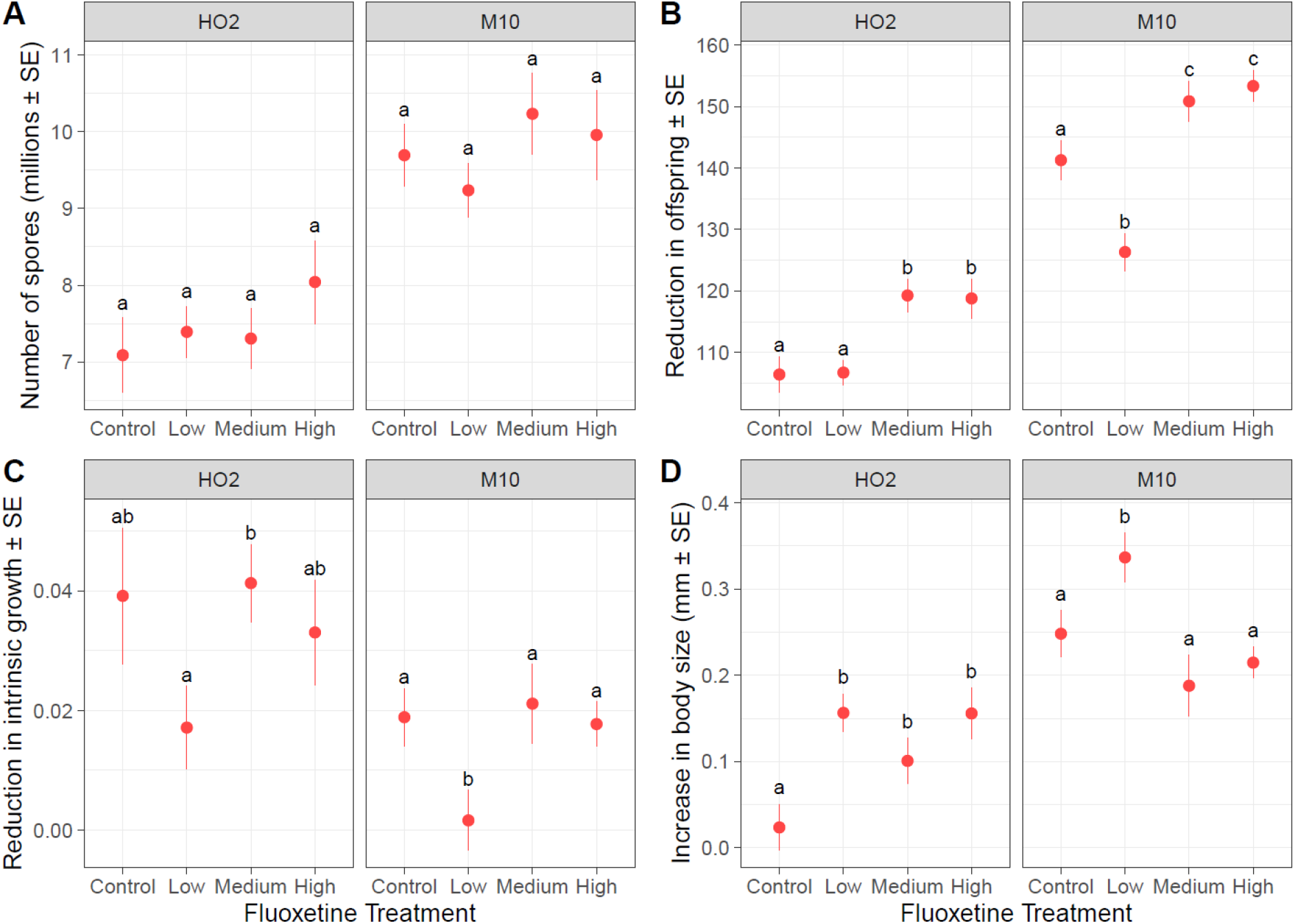
The effect of fluoxetine exposure (control = 0ng/L, low = 26ng/L, medium = 198ng/L, high = 1900ng/L) on A) mature spore loads, B) reduction in the number of offspring produced C) reduction in calculated intrinsic growth and D) increase in body size of infected *Daphnia magna* of two genotypes (HO2 and M10). B, C and D were calculated as the difference in these traits (offspring, intrinsic growth and body size) from the mean trait value of unaffected animals of each respective fluoxetine treatment group. Points represent treatment means (± SE). Lowercase letters indicate significant groupings by post-hoc comparisons conducted separately for each host genotype (P < 0.05).

In contrast to the reduction in fecundity, infected hosts exposed to the low fluoxetine treatment instead had a significantly smaller reduction in intrinsic growth compared to the other fluoxetine treatment groups (Figure 1C). This was true for both host genotypes, as we found no interaction between fluoxetine treatment and genotype (Table 1, Figure 1C). Finally, the change in body size was influenced by an interaction between fluoxetine exposure and genotype (Table 1). Specifically, for infected HO2 *D. magna*, all fluoxetine treatments resulted in a significantly larger increase in body size compared to controls, whereas for M10, only *D. magna* exposed to the low fluoxetine treatment had a larger increase in body size (Figure 1D). Despite the variation in responses to fluoxetine exposure between measures of fitness components and host genotypes, in almost all cases, the greatest magnitude of change occurred at the lowest fluoxetine exposure (the one exception being HO2 reduction in fecundity).

### Fluoxetine exposure alters the relationship between pathogen replication and infection cost in a genotype-specific manner

Our above results show that pathogen spore loads remained relatively constant across different fluoxetine levels, but that the fitness costs imposed by the pathogen instead are sensitive to the concentration of fluoxetine that a host encounters. To further explore how fluoxetine exposure influenced the relationship between pathogen replication and cost of infection, we tested how well the changes in fecundity, intrinsic growth, and body size predicted pathogen spore loads (i.e., slopes in a multiple regression), and if these relationships were modified by fluoxetine treatment, host genotype, or a combination of these factors. Our model fitting approach revealed that fluoxetine exposure changes the relationship between pathogen spore loads and the cost of infection for a host. The exact relationship, however, depended on the specific combination of host genotype and fluoxetine concentration, as the model that included all the three-way interaction terms significantly improved the fit compared to the model where relationship varied by host genotype and fluoxetine independently (Table 2).

**Table 2:** Candidate analysis of covariance models describing the effects of host genotype and fluoxetine treatment on the relationship between pathogen spore loads and the costs of infection (measured as the reduction in fecundity, increase in body size and reduction in intrinsic growth relative to the trait means of control animals). The models are listed in order of complexity, beginning with a null model where the fitness costs are assumed to be unrelated to pathogen spore loads (model 1), and ending with the most complex model where regression slopes varied for every combination of host and fluoxetine treatments (model 6). A log-likelihood ratio test was used to test if the addition of more complex terms improved the fit of the underlying reduced model.

| Candidate models | Terms added | Base model | $\chi^2$ | DF | P-value |
| --- | --- | --- | --- | --- | --- |
| 1. No slopes | Treatment intercepts | - | - | - | - |
| 2. Common slopes for all treatments | Δfecundity<br>Δbody size<br>Δintrinsic growth | 1 | 49.813 | 3 | **<0.001** |
| 3. Different slopes for host genotype only | Host × Δfecundity<br>Host × Δbody size<br>Host × Δintrinsic growth | 2 | 8.755 | 3 | **0.033** |
| 4. Different slopes for fluoxetine levels only | Fluoxetine × Δfecundity<br>Fluoxetine × Δbody size<br>Fluoxetine × Δintrinsic growth | 2 | 13.373 | 9 | 0.146 |
| 5. Different slopes for host genotype and fluoxetine levels independently | Host × Δfecundity<br>Host × Δbody size<br>Host × Δintrinsic growth<br>Fluoxetine × Δfecundity<br>Fluoxetine × Δbody size<br>Fluoxetine × Δintrinsic growth | 2 | 22.564 | 12 | **0.032** |
| 6. Different slopes for every host genotype and fluoxetine levels combination | Fluoxetine × Host × Δfecundity<br>Fluoxetine × Host × Δbody size<br>Fluoxetine × Host × Δintrinsic growth | 5 | 20.002 | 9 | **0.018** |

Examining the regression coefficients estimated separately for each treatment combination (Figure 2) revealed that exposure to fluoxetine never resulted in a significant change in sign for any of the regression coefficients, regardless of host genotype or fluoxetine concentration. For all fluoxetine treatments and host genotypes, spore loads were positively associated with reduction in fecundity and increase in body size, but negatively associated with reduction in intrinsic growth. Instead, fluoxetine exposure modified the strength of these positive and negative relationships in a genotype specific manner. For HO2 hosts, the high fluoxetine exposure resulted in the largest shifts in the slope of the regressions. For M10 hosts, on the other hand, the shifts in the regression slopes were milder overall, while the low and medium fluoxetine exposures were responsible for the largest shifts.

**Figure 2.**
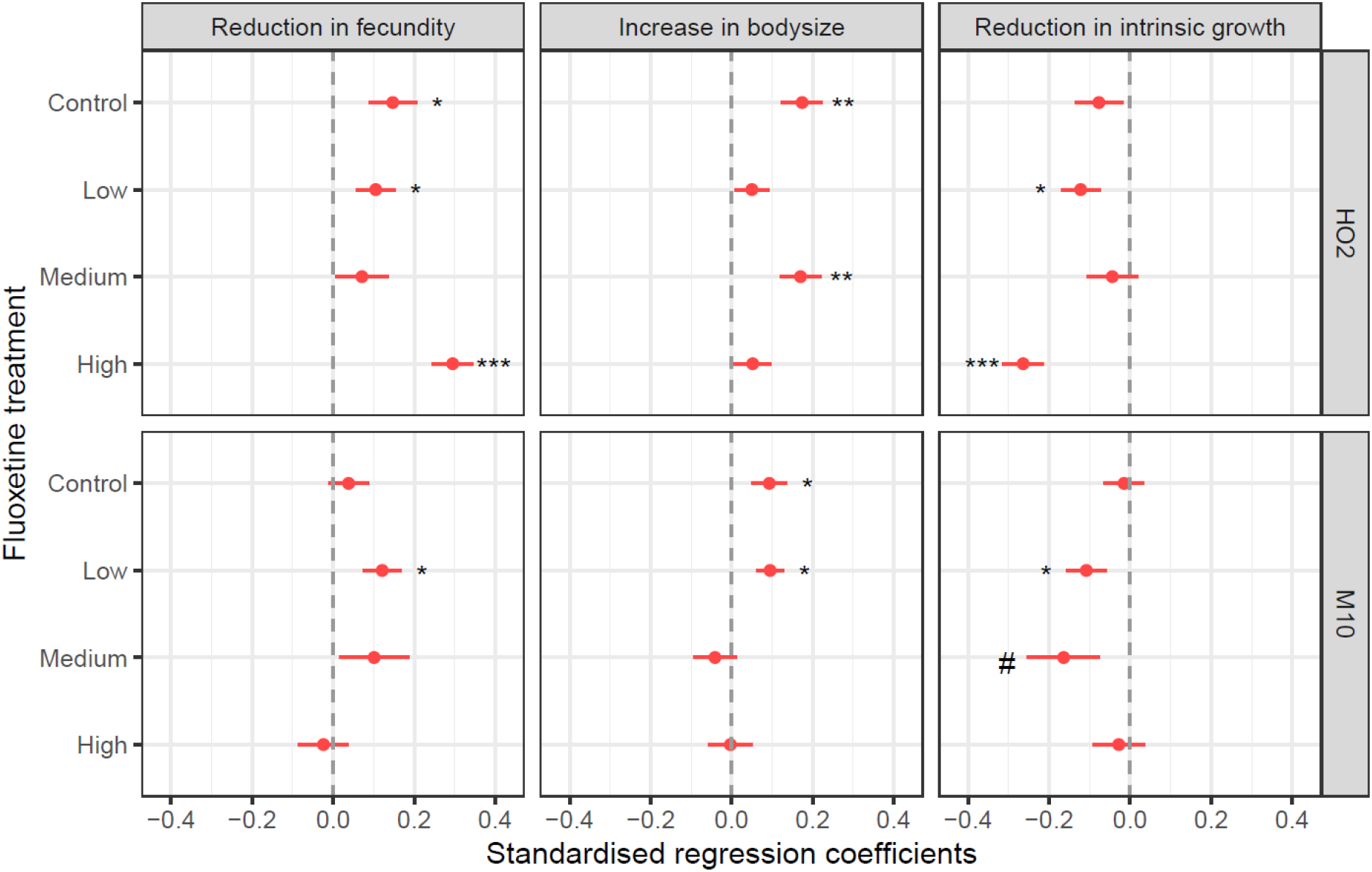
Estimated regression coefficients (± SE) for the association between pathogen spore loads and the standardised reduction in fecundity, increase in body size and reduction in intrinsic growth for hosts of two genotypes (HO2 and M10) exposed to one of four fluoxetine treatments (control = 0ng/L, low = 26ng/L, medium = 198ng/L, high = 1900ng/L). Regression coefficients were estimated separately for each host genotype and fluoxetine treatment combination and spore loads were scaled to a mean of one before analysis. Significance is denoted as * = P <0.05, ** = P <0.01, *** = P <0.001. Marginal significance (P <0.1) is denoted with #.

## Discussion

The costs of infection are viewed as an inevitable consequence of a pathogen’s replication within its host ([4], reviewed in [9]), but the strength of this relationship has been found to be heavily sensitive to environmental conditions (e.g. [13, 14]), including pollutants [13, 18, 21]. A common assumption of many ecotoxicology studies is that the greater the dose of a pollutant, the larger the negative effect [65] – which has indeed been the case in many studies investigating how conventional pollution affects disease dynamics (e.g. [13, 19, 66, 67]). For many of the fitness costs measured here, however, we instead found that the lowest, rather than the highest, fluoxetine concentration resulted in the largest shifts. For example, exposure to just 30ng/L of fluoxetine resulted in infected individuals showing a much milder reduction in intrinsic growth than the control group, while animals exposed to the higher concentrations of fluoxetine experienced the same reduction in intrinsic group as the controls (Figure 1C). While similar non-monotonic effects have frequently been reported on behaviour and life-history traits (e.g. [36, 38, 39, 47]), and in one case pathogen transmission [31], our results demonstrate that the non-monotonic responses typically induced by pharmaceutical pollutants extend to the reduction in fitness that is caused by a pathogen.

Changes in the costs of infection have previously been reported for other pollutants, such as pesticides [18, 68] and heavy metals [19, 20], and it is usually assumed that a corresponding shift in pathogen replication, and ultimately transmission, will occur [18]. In contrast, our results show that the influence of fluoxetine pollution is overwhelming felt though the fitness costs imposed by the pathogen alone, as pathogen load remained largely robust to changes in fluoxetine concentration (Figures 1B–D versus 1A). At least for pharmaceutical pollutants, shifts in the fitness cost of infection do not necessarily result in an equivalent change in pathogen replication, and thus these pollutants may be altering the fundamental trade-off between virulence and pathogen replication (e.g. Figure 2). This could arise if fluoxetine is either acting directly on the way the pathogen is exploiting host resources, or instead modifying the tolerance of the host and its ability to mitigate the damage caused by a pathogen [17, 21].

Regardless of the underlying mechanism, pathogens on average produced more spores when there were relatively larger reductions in fecundity and greater increases in body size (Figure 2), likely because a greater suppression of reproduction and larger body size results in more resources for a pathogen to exploit [18, 69]. Pathogens also produced more spores when the relative reduction in intrinsic growth was smallest (Figure 2). The exact mechanism for this relationship remains unclear, but it suggests that while pathogens perform best when total fecundity is reduced, earlier reproduction, on which intrinsic growth heavily depends, may be favourable for pathogens (see [70]). Fluoxetine modified these relationships, however, in a manner that was entirely dependent on the exposure concentration and the genotype of the host (Table 2). For some fluoxetine and host genotype combinations, variation in spore loads was most strongly associated with different components of virulence (e.g. medium versus high fluoxetine for host HO2, Figure 2). For others, similar fitness costs were linked to spore production, but the relative strengths of each association varied by host genotype or fluoxetine concentration (e.g. variation the reduction in fecundity regression coefficients across all combinations, Figure 2).

Our results indicated that fluoxetine did not alter the direction (i.e., sign) of the relationships between spore loads and the different costs of infection, but instead modulated the strength of any association in a genotype specific manner. This transient nature of the relationship between within pathogen replication and the changes in host fitness helps explain why it is has been difficult to detect clear and consistent virulence-transmission trade-offs experimentally [2, 9], as other factors present, such as pollutants, can modify which fitness components are most affected as well as the strength of the relationship (e.g. [13, 21]). Indeed, our results suggest that pharmaceutical pollution may be a particular powerful modifier the relationship between host and pathogen fitness, as the greatest change, both in terms of the reduction in host fitness, and its relationship to pathogen load (Figures 1 and 2), can arise at the lowest concentration, which represents commonly detected levels in natural populations.

Overall, our study demonstrates how pharmaceutical pollution can influence the relationship between host and pathogen fitness by inducing non-monotonic changes in the damage a pathogen causes a host during infection. This result reiterates how testing unrealistically high concentrations of pharmaceuticals, as is common in the field of ecotoxicology (see discussions in [31, 36, 47]), may distort our understanding of the impact of these emerging pollutants in the wild. It also shows that even ecosystems exposed to trace amounts of pharmaceuticals have the potential to be substantially impacted (see also [31]), highlighting the important role that pharmaceuticals such as fluoxetine may play in shaping the evolution and ecology of host-pathogen interactions in an increasingly polluted world.

## Acknowledgements

We would like to thank Isobel Booksmythe for assisting laboratory work, David Williams and Envirolab Services for analytical testing of water samples, Tobias Hector for assistance with analysis, and Jason Rohr for providing valuable feedback on an earlier version of this manuscript.

