## supplementary material for "Pharmaceutical pollution alters the cost of bacterial infection and its relationship to pathogen load"

Table S1. Measured concentrations of fluoxetine based off water analysis performed by Envirolab Services (MPL Laboratories; NATA accreditation: 2901; accredited for compliance with ISO/IEC: 17025). At each water change, 40mL water samples were taken stored at -18°C until the termination of the experiment, whereupon a subset of these samples were shipped to Envirolab premises, while the remaining samples were kept in storage. Analysis was performed using gas chromatography-tandem mass spectrometry (7000C Triple Quadrupole GC-MS/MS, Agilent Technologies, Delaware, USA) following methods described in [39].

| **Date Sampled** | **Date Analysed** | **Nominal Concentration (ng/L)** | **Measured Concentration (ng/L)** |
|---|---|---|---|
| 08/03/2020 | 28/04/2020 | 30 | 17.2 |
| 08/03/2020 | 28/04/2020 | 30 | 15.8 |
| 08/03/2020 | 28/04/2020 | 3000 | 1500 |
| 10/03/2020 | 28/04/2020 | 300 | 180 |
| 10/03/2020 | 28/04/2020 | 3000 | 1800 |
| 12/03/2020 | 28/04/2020 | 30 | 34.9 |
| 17/03/2020 | 28/04/2020 | 30 | 29.4 |
| 17/03/2020 | 28/04/2020 | 3000 | 1800 |
| 19/03/2020 | 28/04/2020 | 30 | 26.9 |
| 19/03/2020 | 28/04/2020 | 30 | 25.3 |
| 19/03/2020 | 28/04/2020 | 300 | 210 |
| 26/03/2020 | 28/04/2020 | 0 | <2 |
| 26/03/2020 | 28/04/2020 | 30 | 23.3 |
| 26/03/2020 | 28/04/2020 | 300 | 220 |
| 26/03/2020 | 28/04/2020 | 3000 | 2600 |
| 30/03/2020 | 28/04/2020 | 30 | 23.5 |
| 02/04/2020 | 28/04/2020 | 300 | 180 |
| 02/04/2020 | 28/04/2020 | 3000 | 1800 |
